# Vaccination reshapes the virus-specific T cell repertoire in unexposed adults

**DOI:** 10.1101/2020.10.01.322958

**Authors:** Yi-Gen Pan, Benjamas Aiamkitsumrit, Laurent Bartolo, Yifeng Wang, Criswell Lavery, Adam Marc, Patrick V. Holec, C. Garrett Rappazzo, Theresa Eilola, Phyllis A. Gimotty, Scott E. Hensley, Rustom Antia, Veronika I. Zarnitsyna, Michael E. Birnbaum, Laura F. Su

## Abstract

How baseline T cell characteristics impact human T cell responses to novel pathogens remains unknown. Here, we address this question by studying the CD4^+^ T cell response in unexposed individuals to live attenuated yellow fever virus (YFV) vaccine. We quantified virus-specific population dynamics over time using class II peptide-MHC tetramers. Our data revealed that, even in the absence of known viral exposure, memory phenotype T cells were found in the majority of virus-specific precursors in healthy adults. Pre-existing memory T cells can be divided into two groups; abundant pre-vaccine populations that underwent limited overall expansion and rare cells that generated naïve-like responses and preferentially contributed to the memory repertoire after vaccination. Single cell T cell receptor (TCR) sequencing was used to track the evolution of immune responses to different epitopes and showed an association between preservation of unexpanded TCRs before exposure and the robustness of post-vaccine responses. Instead of a further increase in pre-established TCR clones, vaccination boosted the representation of rare TCRs. Thus, vaccine restructures the abundance and clonal hierarchy of virus-specific T cells. Our results link T cell precursor states to post-exposure responses, identifying peripheral education of virus-specific repertoire as a key component of effective vaccination.

∘ YFV-specific precursors contain abundant memory phenotype cells in healthy adults.
∘ Precursor frequency and phenotype are linked to post-immune response.
∘ Vaccination recruits rare virus-specific populations.
∘ Vaccination overrides pre-established clonal hierarchy

## Introduction

The development of a functional memory response is required for protection against subsequent infections (Kathryn et al., 2013; Martin et al., 2012a). Decades of research inspired by questions on immunological memory have led to a wealth of knowledge into T cell responses and the differentiation process that governs memory cell selection (Malherbe et al., 2004; Matthew et al., 2008; Zehn et al., 2009). However, as the majority of studies have used transferred transgenic T cells in laboratory mice, these data have been constrained by cells expressing a limited number of TCR specificities and experimental designs that relied on transferred cells. Using a sensitive peptide-MHC (pMHC) tetramer enrichment protocol that enabled the detection of rare antigen-specific T cells, the endogenous murine responses to antigen challenge were tracked for the first time by Moon et al and Obar et al (Moon et al., 2007; Obar et al., 2008). These seminal studies identified precursor frequency as a key determinant of response kinetics, immunodominance, and memory differentiation. Given the many differences between human and mice, it remains unclear if the same rules apply to human T cell responses to a novel pathogen. This question has gained a new sense of urgency in light of the current pandemic, where several studies have now identified T cells to SARS-CoV-2 in unexposed individuals (Grifoni et al., 2020; Le Bert et al., 2020; Mateus et al., 2020; Peng et al., 2020). Understanding how the composition of the pre-immune repertoire governs post-challenge dynamics will provide the insights necessary for optimizing T cell responses to SARS-CoV-2 and other novel pathogens.

The differentiation state of precursor T cells has generated considerable interest following the identification of SARS-CoV-2-specific memory cells in unexposed individuals (Grifoni et al., 2020; Le Bert et al., 2020; Mateus et al., 2020; Peng et al., 2020). While pre-existing memory T cells to SARS-CoV-2 may be linked to past coronavirus infections, immune memory to unexposed antigens may be a more general phenomenon and has been found in other virus-specific T cells from uninfected adults (Su et al., 2013). Pre-existing memory T cells to the human immunodeficiency virus (HIV) were shown to express the typical memory markers, be capable of rapid cytokine production in vitro, and exhibit evidence of clonal expansion by T cell receptor (TCR) sequencing. Furthermore, HIV-reactive T cell clones from unexposed individuals cross-recognized unrelated microbial peptides, suggesting that pre-existing memory cells may have developed as a cross-reactive response to a broad range of environmental antigens (Su et al., 2013). In mice, a free-living environment has been shown to promote the accumulation of memory T cells and protection against infectious pathogens (Beura et al., 2016; Le Bert et al., 2020; Rosshart et al., 2017). Along with the prevailing paradigm that memory cells provide faster and more robust immune response (Kathryn et al., 2013; Pihlgren et al., 1996; Rogers et al., 2000; Veiga-Fernandes et al., 2000), these findings have generated interest in the possibility that pre-existing memory T cells in an individual could be a mechanism for protection against unencountered pathogens.

In this study, we examined the precursor CD4^+^ T cell repertoire and addressed how precursor states are related to post-vaccination T cell dynamics in humans. We used a highly effective live attenuated yellow fever virus (YFV) vaccine as a model for eliciting primary CD4^+^ T cell response to a novel pathogen challenge. Because YFV does not circulate in the United States and elicits long-lasting antibody responses, individuals naïve to YFV were identified by the absence of exposure and confirmed by a negative serologic test. Longitudinal tracking of CD4^+^ T cell response to YFV was performed using direct ex vivo pMHC tetramer staining and included responses to multiple YFV antigens in the same individual. Our data revealed heterogeneous pre-vaccine repertoires in healthy adults. A substantial number of YFV-specific T cells displayed a memory phenotype, indicating that pre-existing memory T cells can form in the absence of closely related circulating viruses. By following the same T cell specificities over time, we identified differences in T cell kinetics that depended on the pre-vaccination precursor frequencies and initial differentiation states. The clonal composition of YFV-specific T cells before and after vaccination was further assessed by single-cell TCR sequencing and revealed recruitment and predominant expansion of rare TCRs after vaccination.

## Results

### The baseline repertoire of virus-specific CD4^+^ T cells

We immunized seven YFV naïve healthy subjects with the YFV vaccine. These individuals had no history of YFV exposure and were confirmed to be YFV seronegative (Table S1). All volunteers received one vaccine dose and had blood taken before vaccination, 6-10 days, 13-15 days, 25-31 days, and 7 to 34 months after vaccination (Fig. 1A). To enable longitudinal analyses of virus-specific T cells using pMHC tetramers, we recombinantly expressed five HLA-DR monomers matched to donor class II HLA-DR alleles, DRA/B1*0301, 0401, 0407, 0701, 1501. An initial set of 117 peptides was selected based on prior studies and in silico prediction (de Melo et al., 2013; James et al., 2013). Peptide binding to the five HLA-DR alleles of interest was confirmed by competition assay. To focus on relevant YFV epitopes recognized by vaccine-induced T cells, peripheral blood mononuclear cells (PBMC) obtained after vaccination from each individual were stimulated with the selected peptide pool and stained with tetramers (Fig 1B). Tetramers that positively stained cultured cells were then selected for direct ex vivo analyses (Table S2 and Fig. S1A). Tetramer staining was coupled with magnetic column-based enrichment to enable enumeration of rare tetramer-labeled T cells in the unprimed repertoire (Fig. 1C and S1B). Cells were co-stained with anti-CD45RO and anti-CCR7 to delineate the baseline differentiation state of virus-specific T cells. In total, we analyzed 36 YFV-specific CD4^+^ populations using 20 unique epitopes for 5 HLA-DRs in 7 individuals. This revealed CD4^+^ precursor frequencies that ranged from 0.15 to 137 cells per million CD4^+^ T cells (Fig. 1D). In agreement with prior studies on pre-existing memory T cells (Su et al., 2013), memory markers were abundantly expressed by YFV-specific T cells before vaccination (Fig. 1E). Collectively, 40.5% of tetramer^+^ precursors expressed a central memory phenotype (CM, CD45RO^+^CCR7^+^), 12.5% expressed a effector memory phenotype (EM, CD45RO^+^CCR7^−^), and 2.5% expressed a TEMRA phenotype (CD45RO^−^CCR7^−^). The remaining 44.5% were comprised of naïve phenotype T cells (CD45RO^−^CCR7^−^). The abundance of pre-existing memory varied between T cells specific for different epitopes within the same individual and correlated with the size of precursor population (Fig. 1F-G). We looked for shared phenotypes between T cell specificities but did not find a consistent pattern. Cells that recognized the same epitope expressed distinct frequencies and differentiation states in different donors (Fig. S1C). These data indicate that humans can call upon highly variable and heterogeneous T cell repertoires in response to a novel viral infection, including a substantial number of T cells that have already acquired memory differentiation and are abundant before antigen exposure.

**Figure 1:**
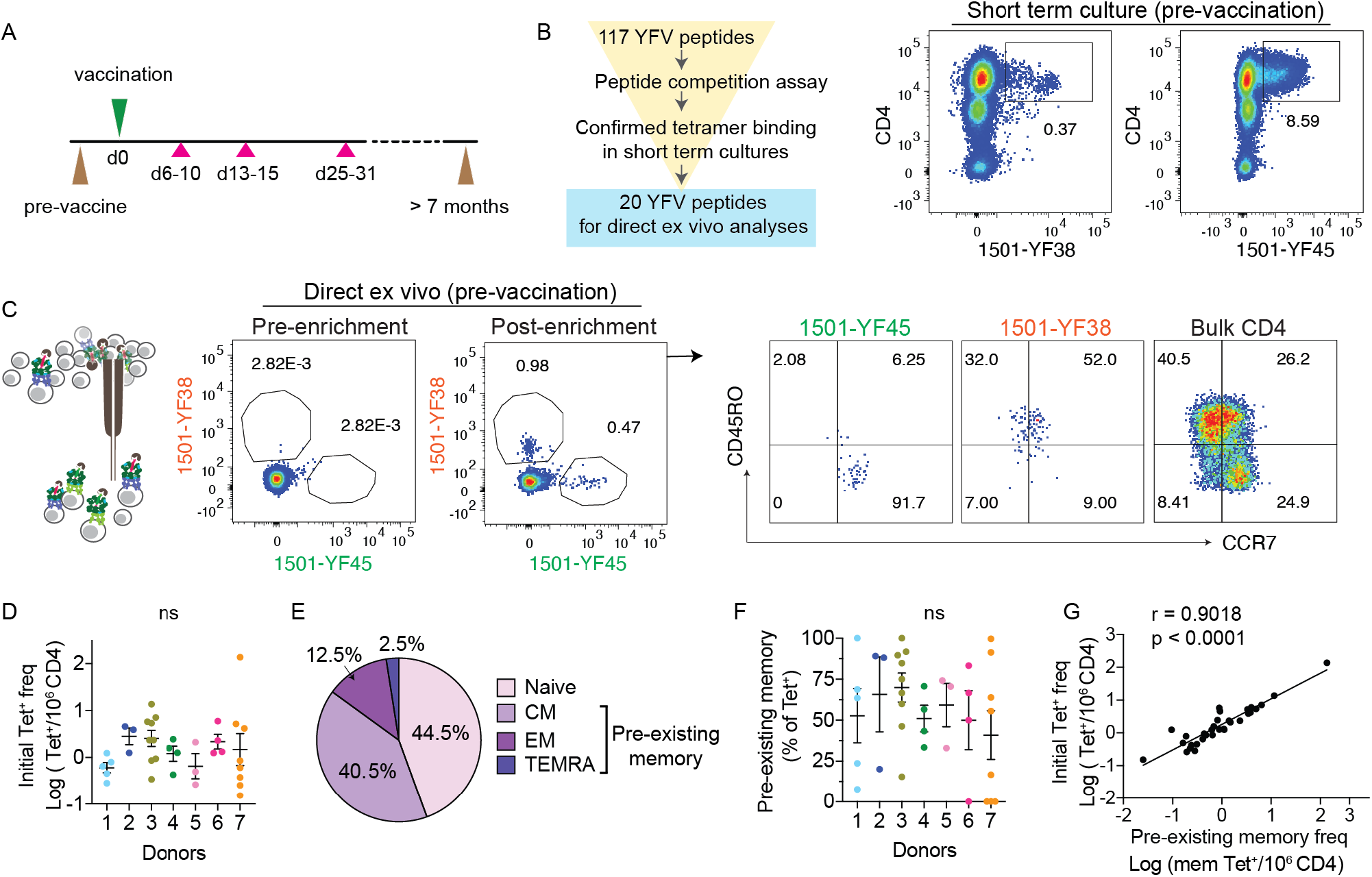
YFV-specific CD4^+^ T cells in YFV unexposed individuals. (A) Overview of the study protocol. At indicated time points, phlebotomy (pink triangles) or leukopheresis (brown triangles) were performed to obtain serial blood samples before and after vaccination. (B) Outline shows the selection process for peptides used in direct ex vivo analyses (left). Example plots show tetramer staining of donor 3 PBMCs obtained on post-vaccine day 14 and stimulated with YFV peptide pool for 3 weeks (right). (C) Direct ex vivo staining, pre- and post-magnetic enrichment, of the same specificities as in B using blood obtained before vaccination. Plots are representative of 4 experiments (left). Anti-CD45RO and CCR7 staining identified tetramer^+^ cells that primarily expressed a naïve (1501-YF45) or memory (1501-YF38) phenotype before vaccination. Bulk CD4^+^ T cells are shown for comparison (right). (D) The frequency of YFV tetramer^+^ CD4^+^ T cell across 7 healthy subjects was quantified by direct ex vivo staining. Each dot represents data from a distinct YFV-specific population, repeated an average of 3.3 times (± 2.3). Donors 1 (n = 5), 2 (n = 3), 3 (n = 9), 4 (n = 4), 5 (n = 3), 6 (n = 4), 7 (n = 8). (E) Differentiation phenotype of tetramer^+^ cells before vaccination (n = 36): naïve (CD45RO^−^CCR7^+^), central memory (CM, CD45RO^+^CCR7^+^), effector memory (EM, CD45RO^+^CCR7^−^), and TEMRA (CD45RO^−^CCR7^−^). (F) Abundance of pre-existing memory T cells as a percentage of tetramer^+^ cells shown in D. (G) Correlation between the frequency of tetramer^+^ T cell and pre-existing memory T cells within each population (n = 36). For D and F, data are shown as mean ± SEM. No statistical difference between donors were identified using Welch’s ANOVA. For G, association was measured by Spearman correlation. Also see Fig. S1.

### Post-vaccine dynamics of primary CD4^+^ T cell response

We next sought to measure the immune dynamics of YFV-specific populations. We used tetramers to monitor vaccine-induced T cell responses during the first month and beyond 7 months after vaccination. Multiple virus-specific populations were tracked simultaneously and identified across time using the same tetramers. This showed that the overall peak of response converged on days 13-15, followed by a typical contraction phase and the establishment of a new baseline at a memory time point (Fig. 2A). To mark activated T cells, tetramer^+^ cells were stained for ICOS, a CD28-related costimulatory molecule induced by TCR signaling (Hutloff et al., 1999). ICOS staining peaked around 2 weeks after vaccination and waned thereafter (Fig. 2B). The incomplete downregulation of ICOS expression a month later suggested that some virus-specific T cells remained activated many weeks post vaccination, possibly in response to antigen persistence after viral clearance (Akondy et al., 2017; Akondy et al., 2009). Quantification of activated T cells using CD38 expression demonstrated a similar pattern (Fig. 2C, S2A). Notably, the summary kinetics do not fully describe the variation in antigen-specific T cell responses. T cells that recognized different YFV epitopes from the same individual generated distinct responses that varied in the rate of expansion, peak frequency, and rate of contraction (Fig. 2D-E). The measured peak frequency differed over 270-fold and ranged between 1.7 to 462 cells per million (Fig. S2B). The maximal effector response generally emerged 2 weeks after vaccination, but a few populations peaked earlier or later and the timing was independent of the initial phenotype (Fig.S2C-D).

**Figure 2:**
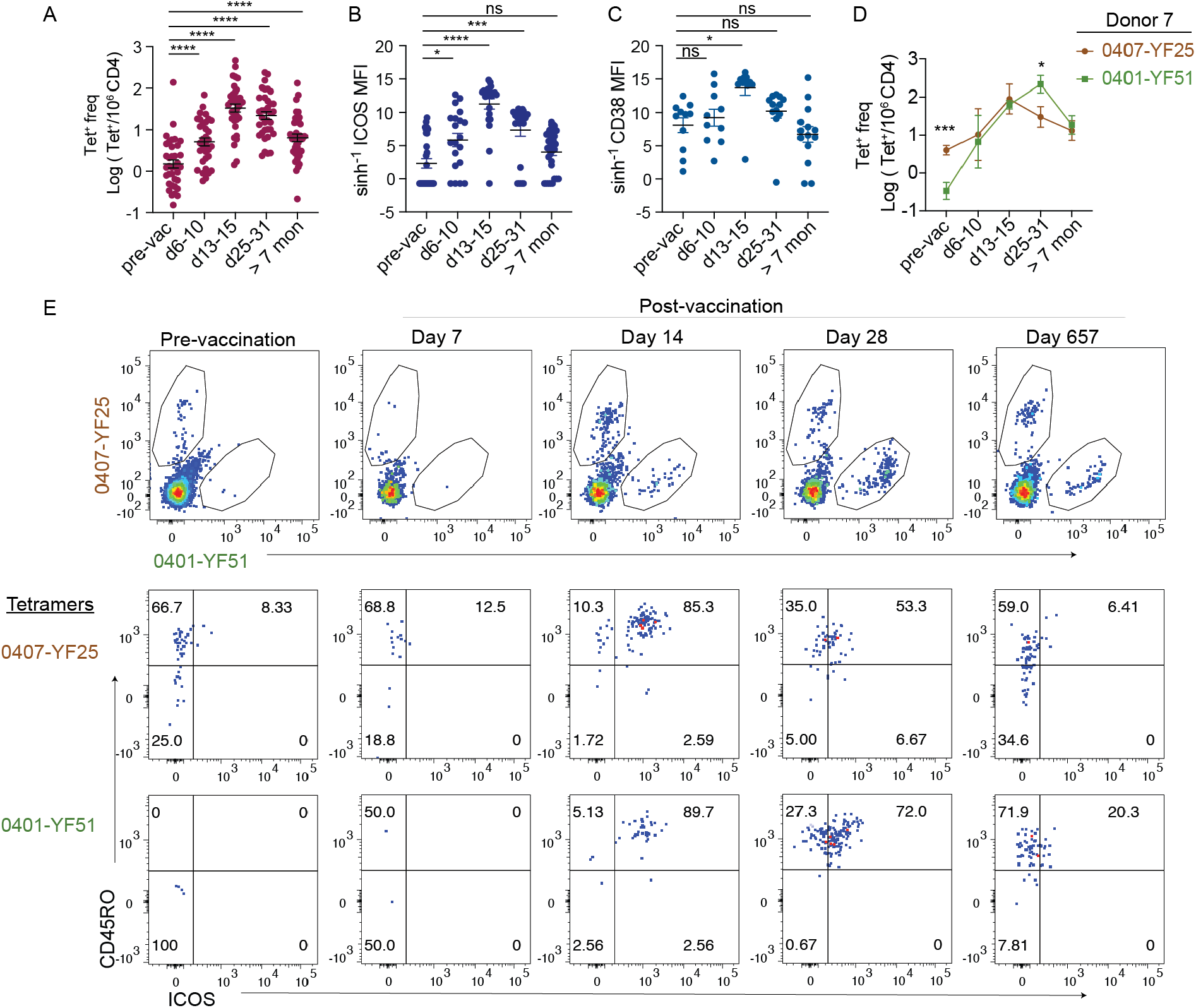
The kinetics of primary CD4^+^ T cell response after vaccination. (A-C) Frequency of tetramer^+^ cells (A) and their activation status as measured by the median fluorescence intensity (MFI) of ICOS (B) or CD38 (C) at the indicated time points (n = 36). (D) Example of two tetramer^+^ populations that showed distinct response kinetics. (E) Plots show changes in tetramer staining for the same populations across time as in D (top) and a time-dependent increase in CD45RO and ICOS expression (bottom). Experiments were performed with the consideration of expected T cell frequency and sample availability (pre-vaccine: 40 million CD4; d7, 14, 28: 10 million PBMCs; d657: 10 million CD4). Plots are representative of 3 experiments. For A-C, mixed-effects analyses were performed. For D, multiple t-test was performed and corrected using Holm-Sidak method. Data are represented as mean ± SEM. * p < 0.05, *** p < 0.001, **** p < 0.0001. Also see Fig. S2.

### Vaccination boosts rare virus-specific populations

Notably, vaccination resulted in a reorganization of virus-specific epitope hierarchies. We compared the frequency of each antigen-specific population months after vaccination with its baseline frequency and found that YFV-specific T cells did not uniformly become more numerous after donors received the immunization. While the vaccine induced an overall increase in YFV-specific T cells, we recovered fewer tetramer^+^ cells for 7 out of 36 virus-specific populations using blood taken 7 months or longer after vaccination (19.4%) (Fig. 3A). We investigated whether early priming events impacted the selection of dominant populations at a later time point. The efficiency by which each population established a memory pool was quantified by dividing the frequency at a memory time point with its starting frequency (> 7months/pre). The magnitude of effector expansion was measured as a fold change between the peak frequency and the pre-vaccine baseline (peak/pre). Comparing these two measurements, we identified a tight correlation between early and late immune responses, such that a boost in the post-vaccine memory frequency was linked to a higher level of effector expansion (Fig. 3B, left). To better understand this relationship, we ranked YFV-specific populations based on the gain in memory cells and compared the temporal dynamics of the top 10 boosted populations with the 7 that lost cells after vaccination. This revealed an intriguing reshuffling of epitope ]hierarchy between the top and the bottom subsets. Populations that shrunk started off at higher baseline frequencies and remained the dominant specificities during the early post-vaccine period, but became surpassed by cells that produced a larger effector response by the second week after vaccination (Fig. 3B, right). Combined data from all donors showed a negative correlation between precursor size and effector response (Fig. 3C). Consistent with this, the rarest precursor subset acquired the largest increase in cell numbers at a memory time point (Fig. 3D). As a group, the relative abundance of T cells that started with an initial frequency of under 1 cell per million represented 3.5% of all tetramer^+^ cells before vaccination and increased to 32.4% after vaccination (Fig. 3E).

**Figure 3:**
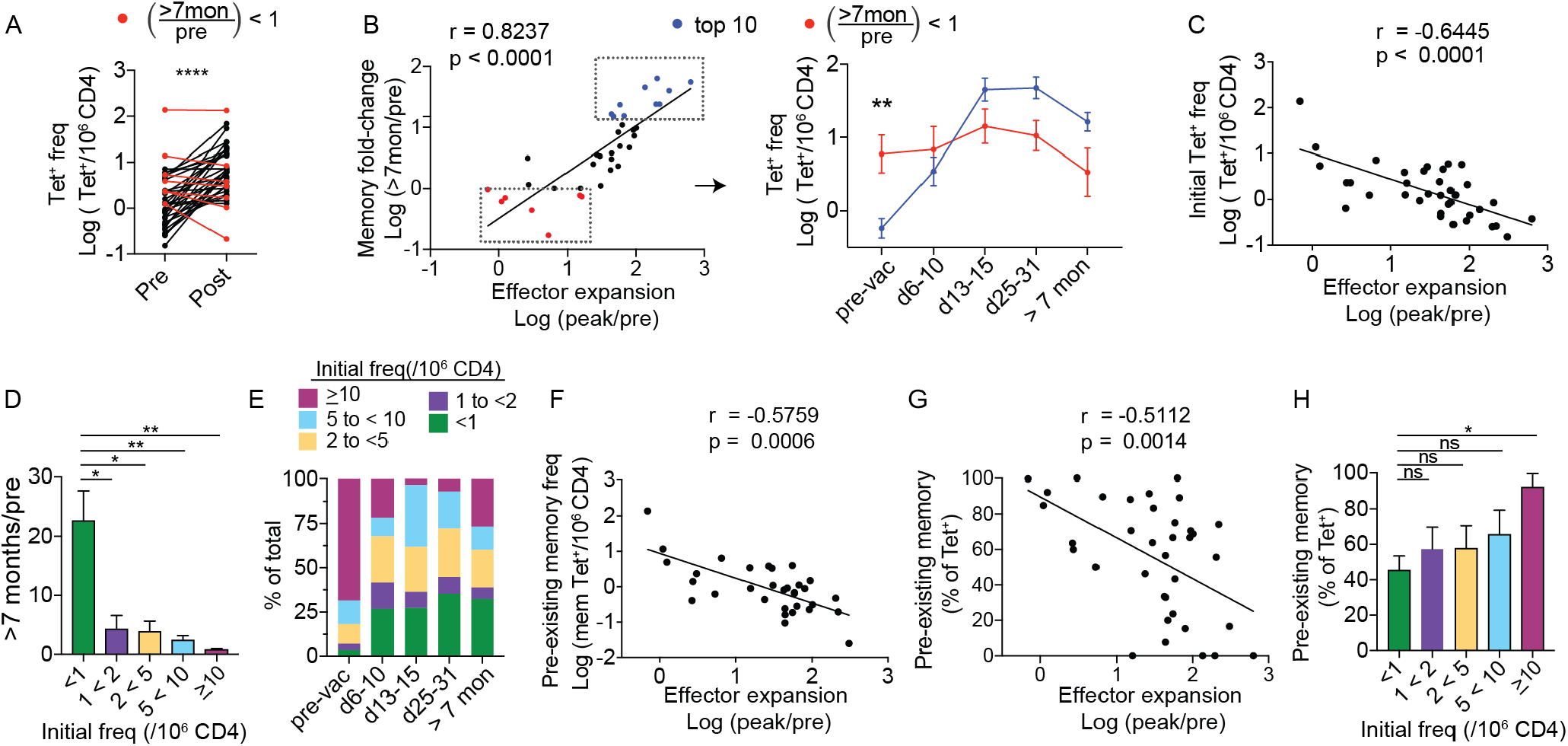
Vaccination preferentially recruits rare tetramer^+^ precursors. (A) Plot show the frequency of tetramer^+^ T cells. Line connects the same population before (pre) and over 7 months after vaccination (post) (n = 36). (B) Correlation between fold-change in effector response and fold-change at a memory time point. Seven populations that became less abundant > 7 months after vaccination are indicated in red. The top ten most expanded populations are indicated in blue. Each dot represents a distinct tetramer^+^ population. Right panel shows longitudinal changes in T cell frequency for the blue and red subsets. (C) Correlation between effector expansion and pre-vaccine frequency. (D) Tetramer^+^ cells are grouped by pre-vaccination frequencies (<1 (n = 15),1 to < 2 (n = 6), 2 to < 5 (n = 8), 5 to < 10 (n = 5), ≥ 10 (n = 2)). Bar-graph shows fold-change in frequency at a post-vaccine memory time point for cells in each bin. (E) The relative abundance of cells in each bin is plotted as a percentage of all tetramer^+^ cells at the indicated time points. (F-G) Correlation between effector response and the frequency (F) or the relative abundance (G) of tetramer^+^ cells that expressed a memory phenotype prior to vaccination. (H) Bar-graph shows the average memory phenotype in each bin as a percentage of tetramer^+^ cells. For A, paired t-test was performed. For B and G, association was measured by Pearson correlation. For C and F, Spearman correlation was used. Differences between blue and red subsets in B were tested by multiple t-test and corrected by Holm-Sidak method. For D and H, Welch’s ANOVA was used. Data are represented as mean ± SEM. * p < 0.05, ** p < 0.01, **** p < 0.0001. Also see Fig. S3.

The low magnitude expansion generated by larger size precursors was unexpected and contrasted with studies in mice, where precursor frequency positively correlated with effector responses (Moon et al., 2007; Nelson et al., 2015). One explanation for this may be the presence of pre-existing memory T cells in the human pre-immune repertoire, which are largely absent among CD4^+^ T cells from adult mice. In support of this idea, the frequency of pre-existing memory T cells inversely correlated with effector response (Fig. 3F). Pre-existing memory T cell abundance, as a percentage of tetramer^+^ population, was also negatively associated with vaccine-induced expansion (Fig. 3G). While memory cells were significantly enriched in the largest and the least responsive precursor subset, the broad distribution of differentiated precursors suggested that other pre-existing memory cells may have the potential to generate more vigorous responses (Fig. 3H). Separating tetramer^+^ cells based on their baseline differentiation states, we found those containing ≥ 50% memory cells gave rise to a wider range of effector responses compared to more naïve-skewed populations (Fig. S3). Thus, precursor differentiation introduces variability into T cell responses, which preferentially recruit rare antigen-specific populations after a primary vaccine challenge.

### Vaccination reshapes the baseline virus-specific repertoire to establish a new clonal hierarchy

We sought to dissect the heterogeneity within pre-existing memory compartment. Because a key goal of vaccination is to recruit virus-specific T cells after exposure, we defined weak responders based on effector responses of the 7 populations that failed to gain cells after vaccination (20, mean peak/pre value plus 2 standard deviation). Tetramer^+^ cells that started with ≥ 50% memory phenotype were partitioned into high (> 20) or low (<20) response groups. Compared with naïve-dominant precursors that contained < 50% memory T cells, low-response populations were significantly more abundant before vaccination (Fig. 4A). Following vaccination, the low-response group was initially more dominant, but became surpassed by other-YFV-specific T cells about 1 week after vaccination (Fig. 4B-C). By contrast, pre-existing memory cells in the high-response group generated a similar kinetic pattern as naïve-enriched precursors. Highly responsive populations, irrespective of the initial phenotype, were less abundant during early post-vaccine period but reached a higher peak frequency and contributed to a higher proportion of memory T cells after vaccination (Fig. 4B-C). The high-response and low-response groups did not differ in their initial differentiation states by CD45RO and CCR7 staining, arguing against differentiation state-related reduction in proliferative capacity (Fig. S4A). The activation kinetics appeared diminished for cells in the low-response subset, although the differences from the other groups were not statistically significant (Fig. S4B).

**Figure 4:**
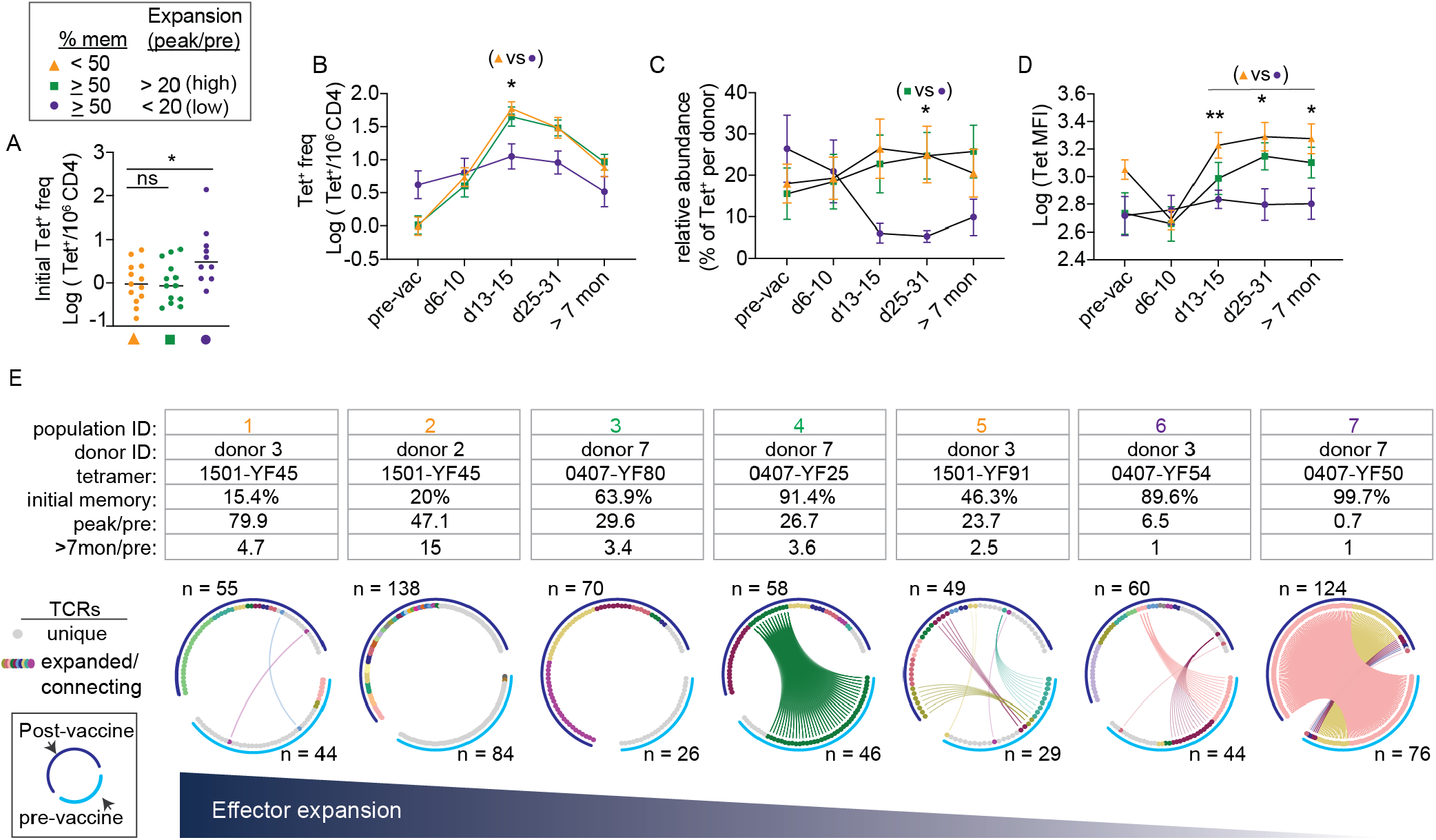
Precursor T cells generate variable responses and undergo repertoire reorganization in response to vaccination. (A-D) Tetramer^+^ T cells were divided into 3 subsets according to the frequency of memory phenotype cells before vaccination and the magnitude of maximal change from the baseline. (A) Scatter plot shows the distribution of pre-vaccination frequency for each subset. (B-C) Change in each subset at the indicated time points by frequency (B) or as a percentage of YFV tetramer-labeled cells identified from the same individual (C). (D) Changes in tetramer staining intensity over time for each subset. (E) Circos plots summarize TCR sequences from 7 YFV-specific populations obtained from 3 donors. Each plot represents a single tetramer^+^ population from one donor and includes data from cells isolated before vaccination (light blue arc) or over 7 months after vaccination (dark blue arc). Gray marks unique TCRs. Other colors indicate expanded TCRs and TCRs found in both time points. TCRs of the same amino acid sequence share the same color, are ordered by frequency, and are connected by lines if found in both visits. For A, Welch’s ANOVA was used. For B – D, mixed effects analysis and Tukey correction was performed. *p < 0.05, **p < 0.01. Data is represented as mean ± SEM. Also see Figures S4-7.

Next we sought to understand how T cells become selected into the post-vaccination repertoire for the subsequent recall response. Past studies using mice have identified affinity-based selection during an immune response. Using tetramer staining intensity to approximate the affinity of TCR-ligand interaction, we observed little change in tetramer MFI after vaccination in the low response group but an increase in other tetramer^+^ T cells, suggesting that T cell expansion was coupled with selective T cell recruitment (Fig. 4D). Consistent with this, the magnitude of effector expansion positively correlated with tetramer MFI at a memory time point after antigen exposure but not before immunization (Fig. S5). To dissect the selection process at a single cell level, we performed TCR sequencing on 903 individually sorted cells encompassing 7 YFV-specific populations from 3 donors using blood drawn before and at least 7 months after vaccination (Fig. 4E, Fig. S6). Each circos plot represented TCRs from the same tetramer^+^ population, shown in separate arcs for the pre-vaccine and post-vaccine time points. TCRs were ordered by frequency and connected by lines if the same TCRβ sequences were shared between blood obtained from separate visits (Fig. 4E, Table S3).

Before vaccination, primarily naïve populations contained mostly unique clonotypes. Expanded TCRs were found in memory-dominant populations, consistent with prior antigen-driven proliferation in the in vivo setting. The degree of clonal expansion was particularly prominent for the most memory-skewed population (population 7), which contained 2 highly expanded clonotypes that occupied 92% of repertoire space. After vaccination, three distinct patterns were observed (Fig. 4E). For populations that started with mostly unique TCR sequences, vaccination focused the immune repertoire onto a diverse set of T cell clones that were minimally shared with TCR sequences from the baseline sample (populations 1, 2, 3). For the most clonal population that generated minimal effector response, the post-immune TCR composition remained remarkably similar to the pre-vaccination repertoire (population 7). For the remaining populations, vaccination established a new clonal hierarchy that replaced the initial repertoire. None of highest ranking clonotypes in the pre-vaccine repertoire remained the most expanded after antigen exposure, although some TCRs detected in the pre-vaccine samples persisted in the post-vaccine blood (populations 4, 5, 6). A clear example of this was population 4, which started with a single dominant expanded clonotype that accounted for 83% of T cells sequenced but became less abundant after vaccination (21%). This clonotype was displaced by a new TCR that was not identified in the baseline sample, suggesting that the T cell(s) expressing the dominant post-vaccine TCR was likely rare before antigen exposure. The recruitment of previously rare T cells into the repertoire reduced the space occupied by pre-expanded TCRs and contributed to a more balanced and diverse clonotype distribution after vaccination (Fig. S7A-B). Notably, vaccination can change the TCR composition of tetramer^+^ cells in the absence of a numerical gain for memory cells after exposure (population 6). Having an inventory of diverse T cells may be necessary to evolve the repertoire, as recruitment of rare precursors failed to occur when the majority of TCRs before vaccination were already expanded (population 7). Measuring the reservoir of singly occurring TCRs as a percentage of total unique clonotypes (diversity reserve), we found a correlation between diversity reserve and the magnitude of early expansion (Fig. S7C). Diversity reserve further correlated with a gain in memory T cells after vaccination (Fig. S7D), suggesting that having a reservoir of rare T cell clones contributed to the expansion and recruitment of virus-specific T cells. Collectively, these data indicate that vaccination reshapes the baseline virus-specific repertoire. For pre-existing memory repertoires that already contain expanded sequences, the presence of alternative TCRs provides the opportunity for recruiting T cells that may be initially rare but are more responsive to viral antigens.

## Discussion

We tracked YFV-specific T cells over time to elucidate characteristics that enabled successful effector response and the recruitment of memory cells. The data described here represent a comprehensive longitudinal study of CD4^+^ T cell response to YFV and include responses to multiple YFV antigens in the same individual. The tetramer-based enrichment approach provided the sensitivity necessary to detect rare precursor T cells and enabled characterization of antigen-specific T cells directly ex vivo in a manner that was not complicated by potential changes from in vitro cultures. Our analyses of the precursor repertoire in YFV naïve individuals showed high level of heterogeneity on population and single cell levels. Tetramer-labeled populations differed over 900-fold in precursor frequency, contained variable abundance of pre-existing memory T cells, and displayed a broad range of TCR diversity. Even though the donors in this study had undergone vigorous screening for YFV exposure, YFV-specific T cells expressing a memory phenotype were identified by tetramers in all individuals before vaccination. These findings are consistent with past studies on other virus-specific memory CD4^+^ T cells in unexposed healthy adults (Su et al., 2013), and together provide strong evidence that pre-existing memory T cells are prevalent in the adult human repertoire. Prior analyses of influenza and HIV tetramer-labeled T cells have identified cross-recognition of other epitopes from unrelated microbes (Su et al., 2013), which was enabled by flexible engagement of TCRs to different pMHC complexes (Reinherz et al., 1999; Reiser et al., 2003). Differences in antigen experience as a result of distinct histories of infection, commensal colonization, and dietary exposures may explain why we observed highly individualized pre-existing memory frequencies among T cells that recognize the same YFV epitope in different people.

A key unresolved question is how pre-existing memory T cells impact human response to novel vaccines and pathogens. Memory cells have generally been viewed a superior source of protective immunity. The advantage of memory cells has been ascribed to their larger starting number and faster division rate (Kathryn et al., 2013; Pihlgren et al., 1996; Rogers et al., 2000; Veiga-Fernandes et al., 2000). However, naïve cells can outcompete memory cells in experiments where low cell numbers or limiting antigens were used, suggesting that the population behavior of memory or naïve T cells is context dependent (Martin et al., 2012b; Mehlhop-Williams and Bevan, 2014). Using pMHC tetramers to track virus-specific T cells over time, we found previously unappreciated heterogeneity within the pre-existing memory repertoire. Cells enriched for pre-existing memory can be divided into subsets that dominated the early or late phases of the immune response. YFV tetramer^+^ cells that started off with higher baseline frequencies were also the most abundant virus-specific T cells in the first week after vaccination. However, they soon became outcompeted by other YFV-specific cells due to a relatively weak capacity to expand. By contrast, the second subset of memory-enriched populations generated naïve-like responses. These cells started off at low frequencies but displayed a robust response to vaccination and preferentially contributed to the memory repertoire after vaccination. These processes lead to a reordered epitope hierarchy that reflect the recruitment of previously rare populations into the memory compartment after vaccination. Our finding that pre-existing memory T cells generate distinct types of responses is highly relevant to the current discussion on SARS-CoV-2-specific T cells in uninfected individuals (Grifoni et al., 2020; Le Bert et al., 2020; Mateus et al., 2020; Peng et al., 2020). The identification of memory T cells that can recognize SARS-CoV-2 before exposure has led to the speculation of pre-existing protective immunity, whereby memory T cells from past coronavirus infections would facilitate a faster and more robust response to SARS-CoV-2 to limit the severity of infection. While we do not know if memory T cells to a never-before-seen virus is qualitatively distinct from those recognizing related virus in circulation, our data would suggest exercising caution in the interpretation of host protection based on the presence of pathogen-specific T cells without additional details on their precursor state and post-vaccine response. Distinct immune dynamics of pre-existing memory may be important for optimizing both early and late T cell responses. Skewing of one subset of pre-existing memory T cells in favor of the other may off-set this balance and promote suboptimal T cell responses to infections.

In addition to reorganizing the immunodominance hierarchy on a population level, vaccination also alters the TCR composition within each virus-specific population. A widely held view is that infectious challenges narrows the TCR repertoire (Busch and Pamer, 1999). Vaccination did indeed selectively expand certain T cell clones from a diverse naïve precursor repertoire. However, pre-existing memory repertoire already contained oligoclonal T cells, which likely have been selected by prior exposures to cross-reactive antigens and may not respond optimally to the current pathogen. We found that the diversity reserve of small clones within a pre-expanded TCR repertoire was associated with the robustness of post-vaccine response. In these populations, vaccination was capable of overriding the pre-established clonal hierarchy to select for rare T cells that were more responsive to vaccine antigens, thereby increasing in the diversity of the expanded clonotypes after vaccination. Having a diverse TCR repertoire is likely beneficial and has been directly linked to protective T cell responses and host survival in mice (Messaoudi et al., 2002). The mechanism of selection may involve cell-intrinsic differences or competition for limited resources such as cytokines and/or antigens. Although our data does not directly address TCR binding characteristics of selected T cells, the finding that tetramer MFI increased after vaccination suggests preferential recruitment of high affinity T cells. Thus, vaccination restructures epitope specificity and clonal hierarchy of virus-specific T cell repertoire. Peripheral education of virus-specific repertoire to cull out the most responsive T cells may be a key mechanism by which effective vaccines generate protective immunity against later pathogen rechallenge.

In conclusion, we have linked precursor repertoire to the primary human CD4^+^ T cell response in the setting of YFV vaccine. Our finding that over half of YFV-specific precursors expressed a memory phenotype suggest that this type of memory cells is not a rare finding in unexposed healthy adults. Different past antigen exposures could impact the ratio of naïve versus memory precursors and expand distinct T cell clones, which will likely influence how T cells from different people respond to a novel infectious challenge. Our data suggest that a critical function of vaccination may be to reshape the baseline repertoire to ensure that the most relevant T cells are recruited for the response to later infection. The impact of advanced age and underlying health conditions on this process may be critical parameters to consider in future designs of effective vaccines against novel pathogens.

## Materials and Methods

### Study subjects

Healthy adults were screened for travel history to YFV endemic regions and prior YFV vaccination by questionnaire. YFV neutralization assay were performed to evaluate antibody response to YFV. All donors were HLA typed (Histogenetics, NY). Seven participants confirmed to have no prior YFV exposure by serologic test and carry the appropriate HLA allele (HLA-DRB1*0301, 0401, 0407, 0701, 1501) were enrolled for the study. Participants received a single dose of 17D live-attenuated yellow fever vaccine strain subcutaneously (YF-VAX^®^, Sanofi Pasteur). Leukopheresis was performed prior to and 210-1011 days after vaccination to obtain sufficient cell numbers. Phlebotomy for whole blood was obtained 6-10 days, 13-15 days, 25-31 days after vaccination. All samples were de-identified and obtained with IRB regulatory approval from the University of Pennsylvania. Subject characteristics are shown in Table S1.

### YFV-17D focus reduction neutralization tests (FRNT)

Approximately 300 foci of YFV-17D were added to two-fold serial dilutions of heat-inactivated serum. Virus-serum mixtures were then incubated for 1 h at 37°C and then added to 2.5E4 Vero cells/well in 96-well flat-bottom plates. Virus-serum mixtures were incubated with cells at 37°C for 1hour, washed, and then overlayed with a 1.5% Methyl Cellulose:1xPa (DMEM) medium. After 40 hours of incubation at 37°C, plates were fixed with 4% paraformaldehyde, permeabilized, blocked, and stained by sequential incubation with a biotin-conjugated 4G2 monoclonal antibody (ATCC HB-112), streptavidin-HRP (BD Biosciences), and TrueBlue peroxidase substrate (KPL). FRNT50 titers were reported as the highest reciprocal dilution giving a focus count ≤ the 50% neutralization cutoff, and the geometric mean was computed for technical duplicates.

### Epitope selection

YFV peptide candidates included sequences identified in previous studies (de Melo et al., 2013; James et al., 2013) and additional peptides predicted to bind HLA-DRB1*0301, 0401, 0407, 0701, 1501 with a consensus percentile under 20 using IEDB analyses resource (Paul et al., 2015). To validate HLA binding, a low stringency peptide competition assay was performed to identify peptides that can compete off biotinylated control peptides by at least 30%. Biotinylated sequences were HA306 (GGPKYVKQNTLKLAT) for DRB1*0401 and 0407, CLIP103 (CGGGPVSKMRMATPLLMQA) for DRB1*0701 and 1501, and TT511 (CGGKIIVDYNLQSK) for DRB1*0301. Biotinylated control peptides were exchanged with HLA-DR at 1:2 molar ratio in the presence of 5 times excess of a test peptide and detected using horseradish peroxidase (HRP)-conjugated streptavidin (BD Biosciences). PBMCs obtained after vaccination from each donor were stimulated with 47 DR-binding peptides at 0.4ug/ml per peptide for 3 weeks and stained with tetramers. For production of tetramers, HIS-tagged DR protein monomers were produced by Hi5 insect cells and purified using Ni-NTA (Qiagen) and size exclusion columns (AKTA, GE Healthcare). Protein biotinylation, peptide exchange, and tetramerization were performed using standard protocols as previously described. (Day et al., 2003; Su et al., 2013). See Table S2 for peptide sequences used for generating the tetramer for direct ex vivo analyses.

### Antigen-specific T cell analyses and sorting

PBMCs were isolated from blood samples through density gradient centrifugation (Ficoll-Paque, GE Healthcare). CD4^+^ or CD3^+^ cells were enriched from leukopheresis products through negative selection (RosetteSep, STEMCELL Technologies) before density gradient centrifugation. Precursor analyses were performed using 30 to 100 million CD3^+^ or CD4^+^ enriched T cells. For post-vaccination analyses, 10 million PBMCs (up to 1 month after vaccination) or 10 to 30 million CD3^+^ or CD4^+^ enriched T cells (> 7 months) were used. Tetramer staining was carried out as previously described (Su et al., 2013). In brief, cells were stained at room temperature for 1 hour using 1.25ug of each tetramer in 50ul reaction. Up to 4 tetramers were combined in the same reaction. Tetramer tagged cells were enriched by adding anti-PE and/or anti-APC magnetic beads and passing the mixture through a magnetized column (Miltenyi). The tetramer-enriched samples were further stained with live/dead dyes, exclusion markers (CD19 and CD11c), anti-CD3, anti-CD4, anti-CD45RO, anti-CCR7, anti-CD38 and anti-ICOS antibodies for 30 minutes at 4°C. Samples were acquired by flow cytometry using LSRII (BD) or sorted on FACS Aria (BD). Frequency calculation was obtained by mixing 1/10^th^ of sample with 200,000 fluorescent beads (Spherotech) for normalization (Su et al., 2013). Data analyses were performed using FlowJo (Tree Star).

### Single-cell TCR sequencing and analyses

Single cell TCR Sequencing by nested PCRs was performed using the primer sets and the protocol as previously described in Han et al. (Han et al., 2014). In brief, reverse transcription was performed with CellsDirect One-Step qRT-PCR kit according to the manufacturer’s instructions (CellsDirect, Invitrogen) using a pool of 5’ TRVB-region specific primers and 3’ C-region primers. The cDNA library was amplified using a second set of multiple internally nested V-region and C-region primers with HotStarTaq DNA polymerase kit (Qiagen). The final PCR reaction was performed on an aliquot of the second reaction using a primer containing common base sequence and a third internally nested Cβ primer. PCR products were gel purified (Qiagen) and sequenced on NextSeq 500/550 (Illumina) by 300 cycle pair-end reaction. Output data (.bcl files) were converted to fastq format using bcl2fastq software. Reads 1 and read 2 were converted into one paired end read using pandaseq (Masella et al., 2012). Data were demultiplexed by the unique combinations of plate, row, and column nucleotide barcodes. Consensus TCRβ sequences were identified using the V(D)J alignment software MiXCR (Bolotin et al., 2015). Successfully aligned sequences were first thresholded by a read count of 200 reads per sequence. In wells containing multiple TCRβ chains above this cutoff, dual chains (two productive chains in a single cell) were called when both chains were separated by a read count ratio of 0.5 or more. In cases for which more than 2 chains passed this criterion or when a TCR sequence was not unique to one tetramer, data were discarded as ambiguous. For downstream analyses, data wrangling was perform using the tidyverse package. TCRs common to pre- and post-vaccine visits were identified using the most abundant TCRβ sequence if more than one sequence was found at a given position. Circos plots were made using the circlize package of R software (Gu et al., 2014).

### Quantification and statistical analyses

Data transformation was performed using Logarithmic function or hyperbolic sine transformation (sinh^−1^ = In 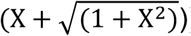) when data contained zero values. Assessment of normality was performed using D’Agostino-Pearson test. Spearman was used if either of the two variables being correlated was non-normal. Otherwise, Pearson was used to measure the degree of association. The best-fitting line was calculated using least squares fit regression. Statistical comparisons were performed using two-tailed Student’s t-test or Wilcoxon signed-rank test, using a p-value of <0.05 as the significance level. Multiple-way comparisons were performed using ANOVA or mixed effect model and corrected for multiple comparisons. Welch’s correction was applied for unequal variance. Statistical analyses were performed using GraphPad Prism. Lines and bars represent mean and variability is represented by standard error of the mean (SEM). * *P* < 0.05, ** *P* < 0.01, *** *P* < 0.001, **** *P* < 0.0001.

## Acknowledgments

We thank Lea Williams, Tammy Ruozhang Xu, and Ning Jiang for helpful discussions.

## Author contributions

Conceptualization, L.F.S.; Experimentation, Y.P., B.A., Y.W, T.E.; Sequence analyses, L.B., P.V.H., and G.C.R; Study recruitment: C.L. and A.M; Modeling and statistical support, R.A., V.Z., P.G.; Supervision, S.H., M.B., L.F.S.; Manuscript preparation, L.F.S. and Y.P.

## Data and materials availability

Data generated or analyzed in this study are included in this article and its supplementary files.

## Supplementary Figures and Tables

**Supplementary Table 1:** Demographic information and neutralizing YFV antibody titers of study participants

| Donor ID | age | sex | Travel history | Nab titer (pre-vaccine) | Nab titer (d13-15) | Nab titer (d25-31) |
| --- | --- | --- | --- | --- | --- | --- |
| 1 | 36 | M | None out of US (not vaccinated) | 1:10 | 1:320 | 1:2560 |
| 2 | 31 | F | None out of US (not vaccinated) | 1:20 | 1:640 | 1:2560 |
| 3 | 63 | M | None out of US (not vaccinated) | 1:10 | 1:320 | 1:2560 |
| 4 | 53 | M | None out of US (not vaccinated) | 1:10 | 1:320 | 1:320 |
| 5 | 27 | M | None out of US (not vaccinated) | 1:40 | 1:1280 | 1:1280 |
| 6 | 24 | M | Venezuela (not vaccinated) | 1:10 | 1:2560 | 1:2560 |
| 7 | 50 | F | Tunisia (not vaccinated) | 1:10 | 1:320 | 1:2560 |

**Supplementary Table 2:**
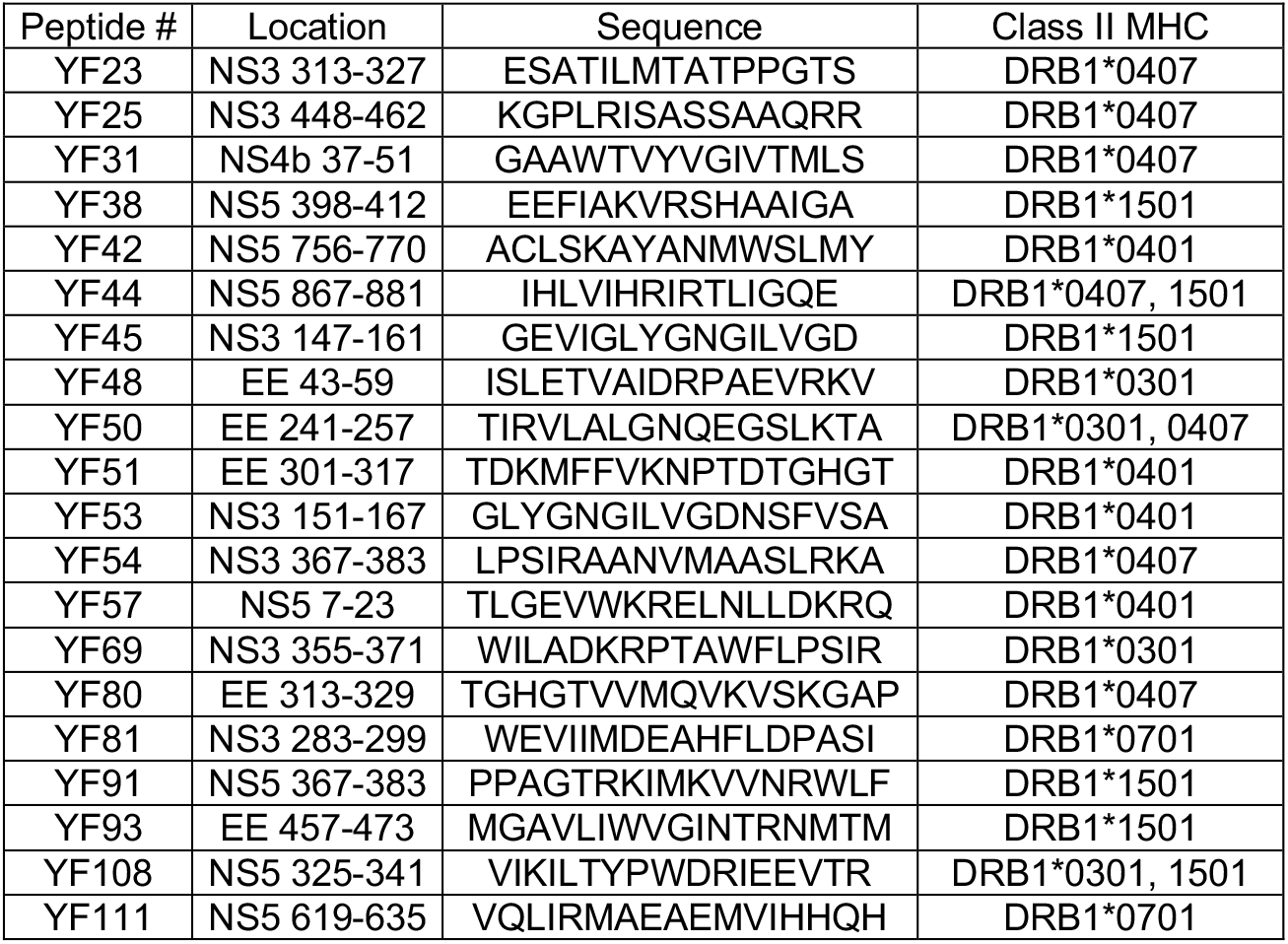
YFV peptide sequences for direct ex vivo analyses

Supplementary Table 3: YFV-specific TCR sequences (see excel spreadsheet)

**Figure S1:**
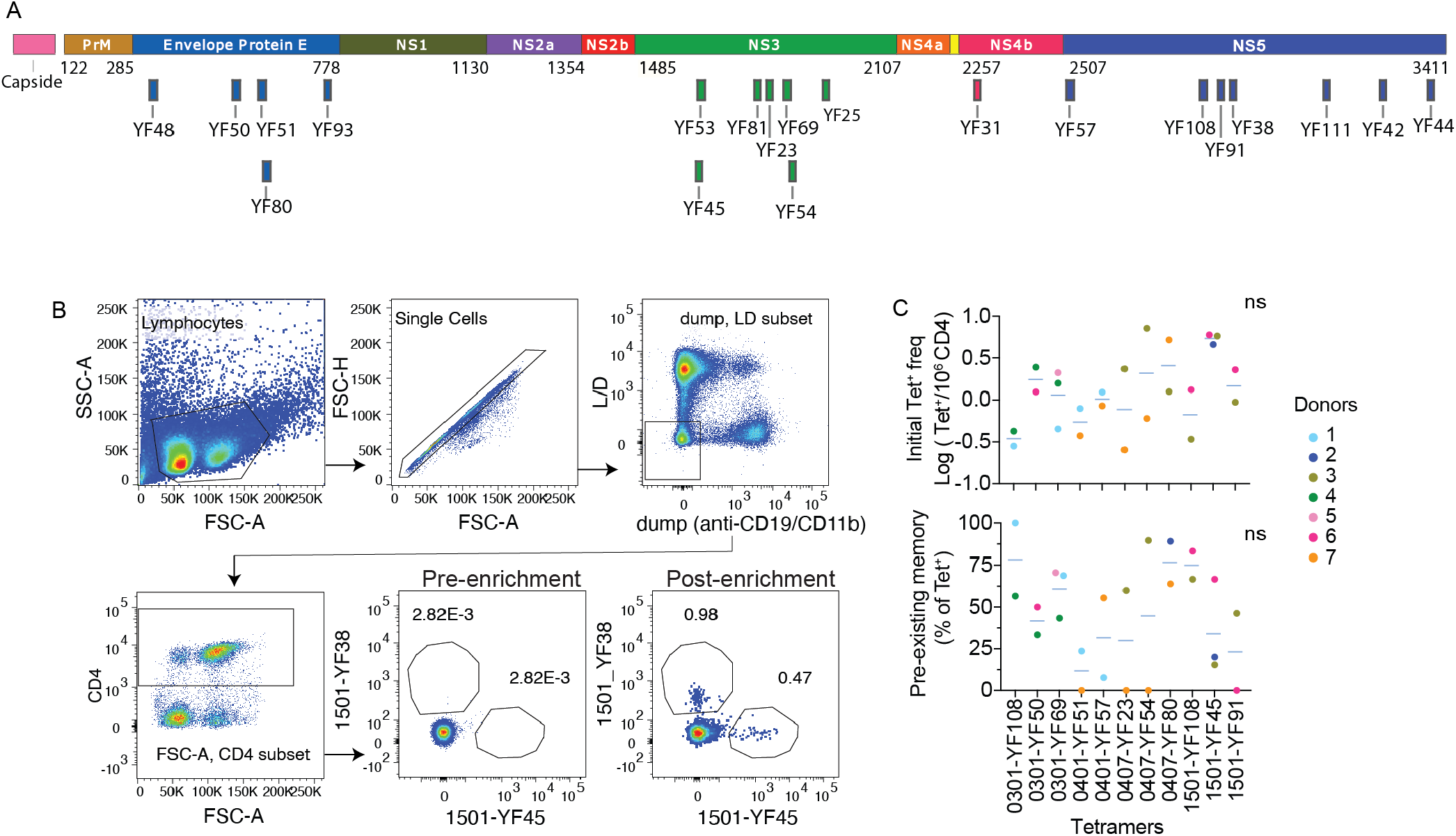
YFV epitopes and T cell analysis across individuals. (A) Epitope map shows the position of 20 YFV peptides used in direct ex vivo analyses. (B) Representative plots show the gating strategy for identifying tetramer^+^ cells shown in Fig. 1C. (C) Scatter plots compare the frequency (top) and memory phenotype (bottom) of T cells that recognize the same pMHC complex in different donors using blood obtained before vaccination. Each dot represents a distinct YFV-specific population and is colored by donor. For C, no statistical difference between tetramer^+^ cells were identified using Welch’s ANOVA. Relates to Figure 1.

**Figure S2:**
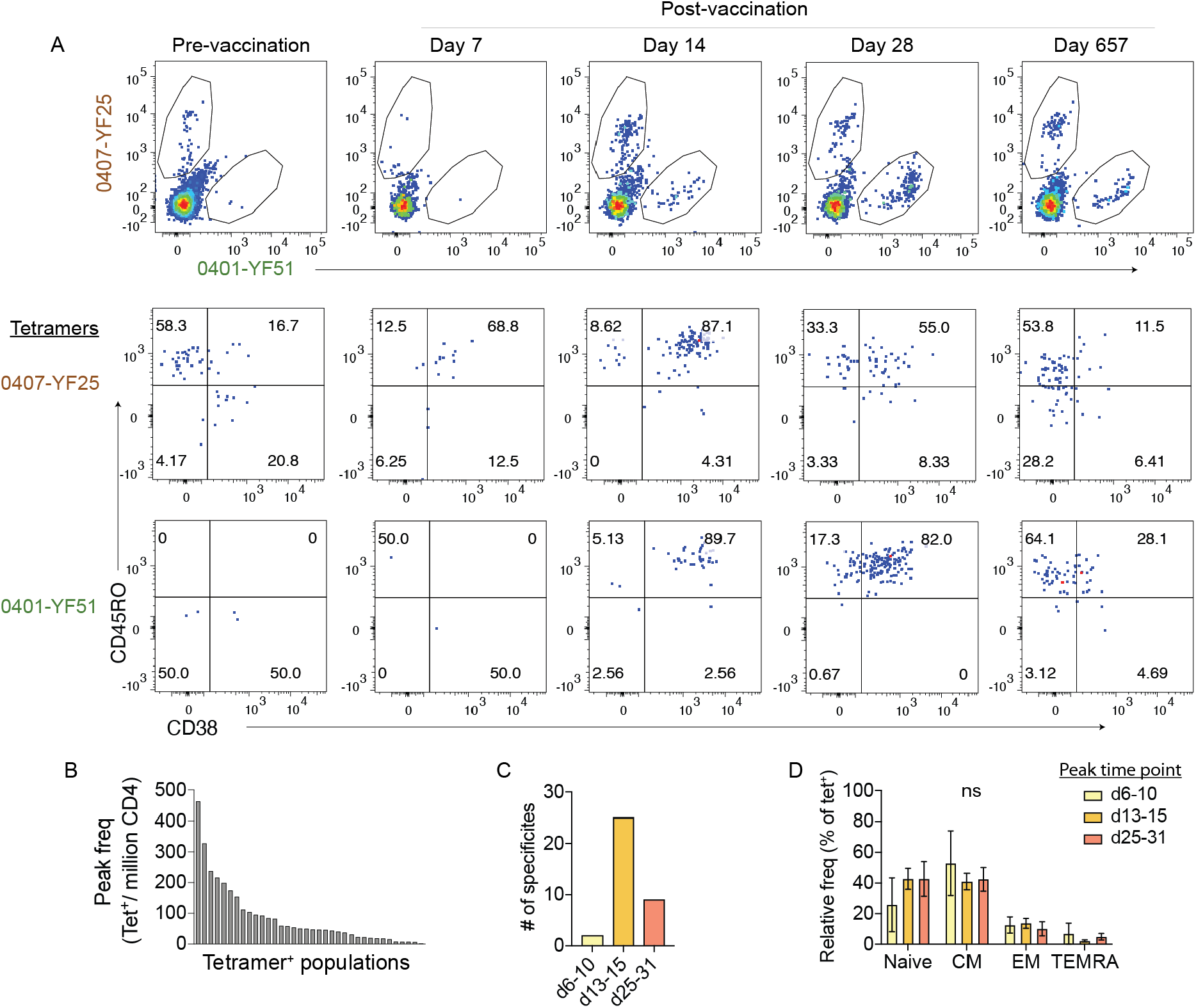
Post-vaccine frequency distribution and activation phenotype. (A) Representative plots show changes in tetramer staining (top) and CD38 expression (bottom) by the indicated tetramer^+^ populations across time points. (B) Plot ranks the maximal post-vaccine frequency achieved by each tetramer^+^ population within the first month after vaccination. (C) Measured maximal frequency occurred on days 6-10 (5.6%, n = 2), days 13-15 (69%, n = 25) or days 25-31 (25%, n = 9) after vaccination. (D) Pre-vaccination phenotype is similar between T cells that peaked at different times. Tetramer^+^ cells were divided based on the timing of maximal effector response. Bar-graph shows the relative abundance of each group expressing a naïve, central memory (CM), effector memory (EM), or TEMRA phenotype before vaccination. Comparison was performed using Welch’s ANOVA. Data are represented as mean ± SEM. Relates to Figure 2.

**Figure S3:**
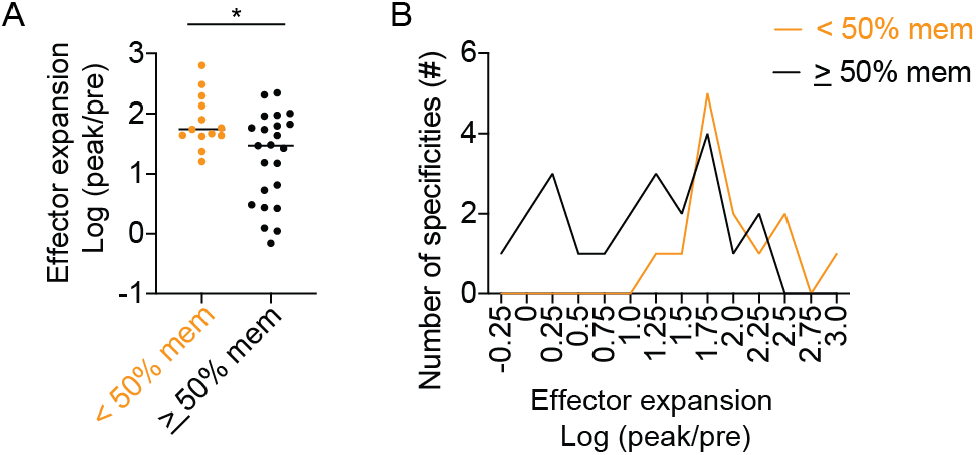
Pre-existing memory T cells generate heterogeneous responses. Tetramer^+^ cells identified in the pre-vaccination samples were divided by the baseline differentiation phenotype into those that contained at least 50% or under 50% of pre-existing memory cells. (A) Scatter plot shows the distribution of effector responses by the indicated subsets. (B) Plot summarizes the number of YFV-specific populations in each group that generated a response within the indicated range. For A, t-test was performed. * p < 0.05. Relates to Figure 3.

**Figure S4:**
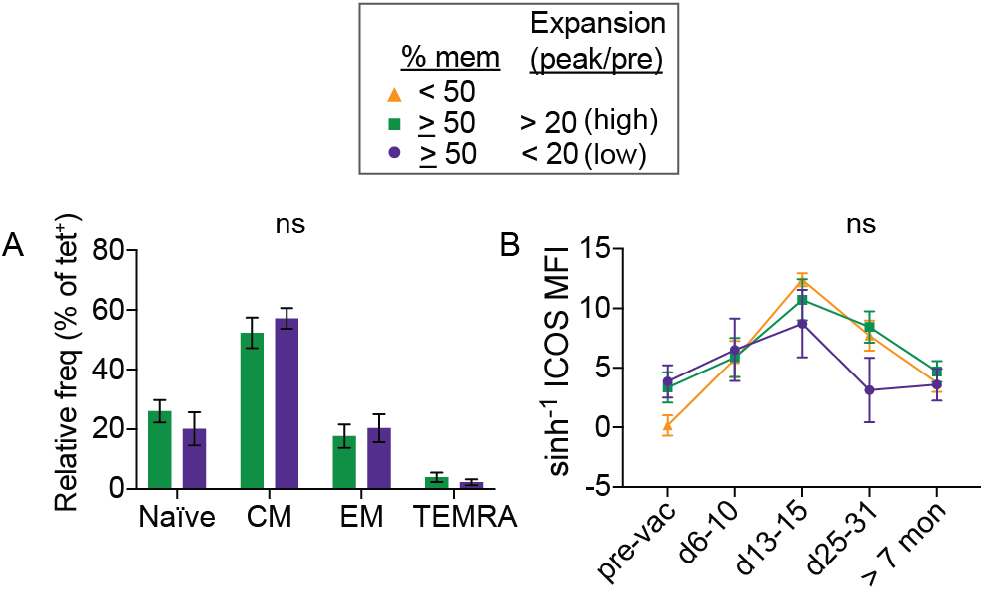
Initial phenotype and activation kinetics of high- and low-response subsets. (A-B) Tetramer^+^ T cells were divided into 3 subsets according to the frequency of memory phenotype cells before vaccination and the magnitude of maximal change from the baseline. Pre-vaccination phenotype is similar between pre-existing memory-enriched populations (memory ≥ 50%) that generated > or < 20-fold-change in effector responses. Bar-graph shows the relative abundance of naïve, CM, EM, or TEMRA subsets within cells in the high- or low-response group before vaccination. (B) Changes in each subset at the indicated time points by ICOS expression. For A, multiple t-test was used and corrected by Holm-Sidak method. For B, mixed effects analysis and Tukey correction were performed. Data are represented as mean ± SEM. Relates to Figure 4.

**Figure S5:**
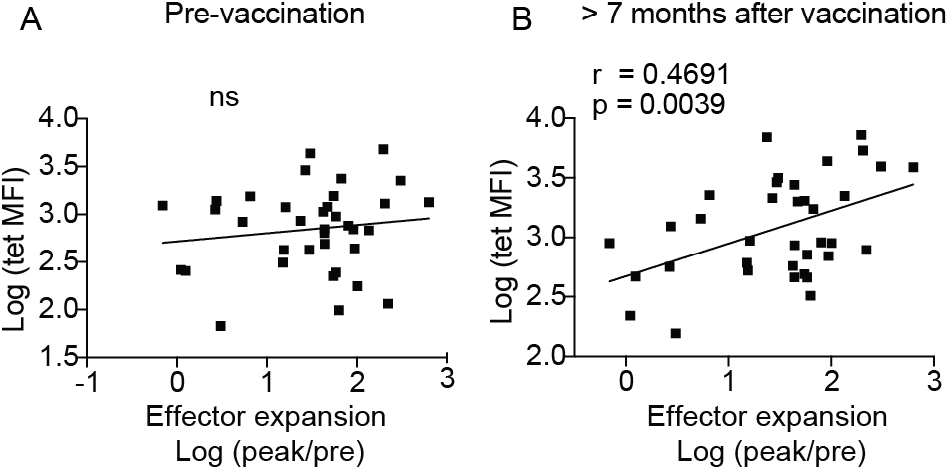
The staining intensity of tetramer^+^ cells increased after vaccination. (A-B) Correlation between fold-change in effector response (peak frequency/pre-vaccine frequency) and tetramer MFI before (A) or > 7 months after vaccination (B). Association was measured by Pearson correlation. Relates to Figure 4.

**Figure S6:**
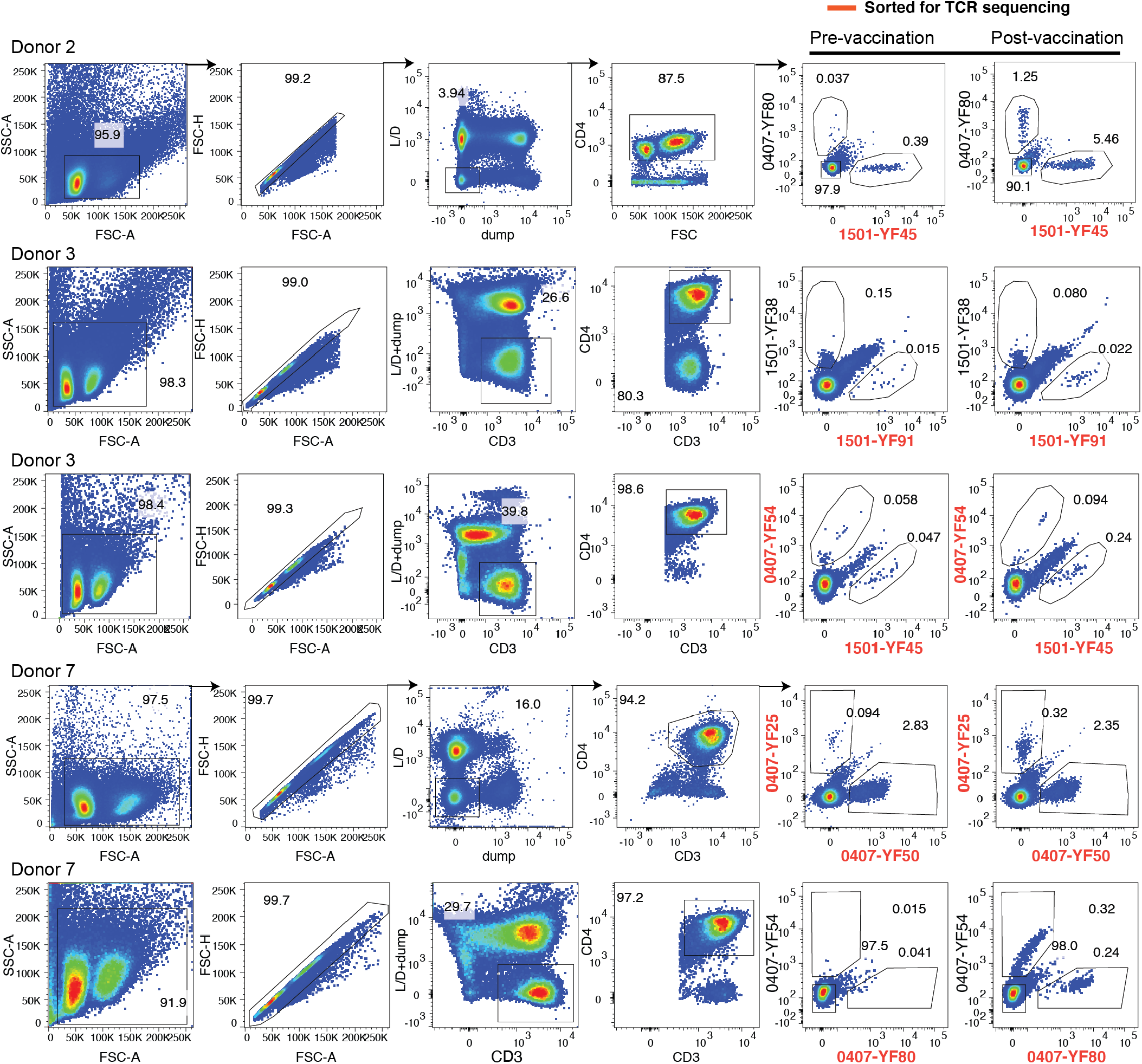
Gating strategy for T cells sorted for TCR sequencing. FACS plots show the sort layout for obtaining tetramer^+^ T cells used for TCR analsyes. Sorted populations are highlighted in orange. Relates to Figure 4.

**Figure S7:**
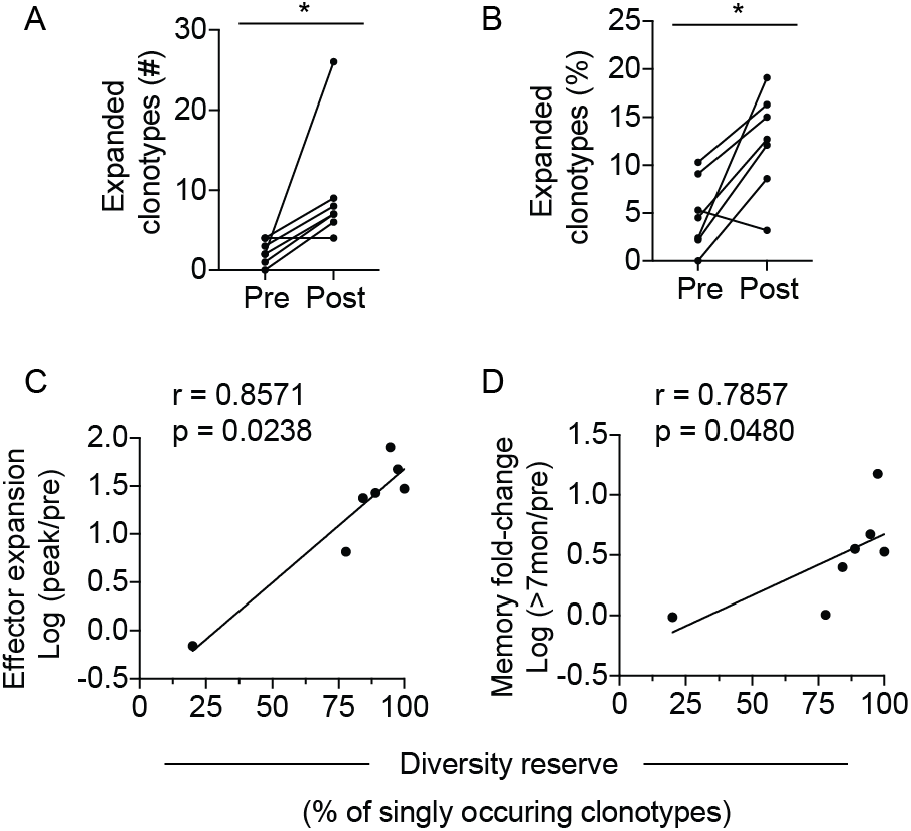
Vaccination response is associated with diversity reserve and an increase in expanded post-vaccine clonotypes. (A) Plot shows the number of distinct clonotypes that are found in 2 or more cells. (B) Frequency of expanded clonotypes as a percentage of the total numbers of TCRs sequenced for each tetramer^+^ population. For A-B, lines connect data from the same tetramer^+^ poulation isolated from blood obtained before vaccination and over 7 months after vaccination. (C) Correlation between the diversity reserve (numbers of single TCRs/numbers of unique clonotypes) and effector response. Diversity reserve was calculated based on tetramer^+^ TCR sequences obtained from pre-vaccination sample. (D) Correlation between diversity reserve and fold change in T cell frequency at a memory time point. For A and B, Wilcoxon matched-pairs signed rank test was used. For C and D, association was measured by Spearman correlation. * p < 0.05. Relates to Figure 4.

